# Characterising recent antimalarial resistance in West Africa: Insights from amplicon sequencing of 17,384 *Plasmodium falciparum* infection samples

**DOI:** 10.64898/2026.02.21.705767

**Authors:** Joyce M. Ngoi, Eniyou C. Oriero, Eyyub S. Unlu, Kukua Thompson, Dzidzor Ayeke, Mona-Liza E. Sakyi, Collins M. Morang’a, Brandon Amambua-Ngwa, Martha Anita Demba, Benjamin Kobna Njie, Fatoumatta Cham, Balla Gibba, Ignatus N. Dorvi, Enock K. Amoako, Charles Mensah, Albert Yao Kudakpo, Oumou Maiga-Ascofare, Abdoulaye Sadio, Thomas Pemberton, Andrew Mains, Mozam Ali, Julia Jeans, Ísla O’Connor, Eleanor Drury, Katherine Figueroa, Matthew Forbes, Antonio Marinho da Silva Neto, Simon Suddaby, Thomas Maddison, Katherine Rowlands, Jordi Landier, Antoine Claessens, Bahdja Boudoua, Issaka Sagara, El-Hadj Ba, Eugene Lama, Tobias O. Apinjoh, Vincent N. Ntui-Njock, Ambroise Ahouidi, Cyrille Diedhiou, Ndeye Khady Sow, Mattu T. Kroma, David J. Conway, Umberto D’Alessandro, Souleymane Mboup, Sónia Gonçalves, Jacob Almagro-Garcia, Kevin Howe, Richard D. Pearson, Cristina V. Ariani, Shavanthi Rajatileka, Victoria J. Simpson, Dominic P. Kwiatkowski, Gordon A. Awandare, Alfred Amambua-Ngwa, Lucas N. Amenga-Etego

**Author notes:** Joint corresponding authors: Alfred Amambua-Ngwa, Lucas N. Amenga-Etego. These authors contributed equally to this work (Joyce M. Ngoi, Eniyou C. Oriero, Eyyub S. Unlu).

## Abstract

*Plasmodium falciparum* (*P. falciparum*) infection remains a significant public health threat in West Africa, where chemoprevention and first-line therapies are key interventions against malaria. However, the development and spread of resistance to commonly used antimalarials poses a growing threat to the efficacy of these strategies. This study characterises the recent landscape of antimalarial resistance in West Africa by analysing targeted amplicon sequences from 17,384 *P. falciparum* infection samples. Across countries, the prevalence of the pyrimethamine resistance–associated *dhfr* triple mutant allele (51I/59R/108N) exceeded 80%, while its combination with the sulphadoxine resistance–associated *dhps* 437G exceeded 60% of infections. Unlike the parasite genotypes in East Africa, the prevalence of the *dhps* 540E mutant was low (1.5%), whereas *dhps* 436A was common (43.8%). The chloroquine resistance marker *crt* 76T showed greatest geographic heterogeneity, ranging from low prevalence in Ghana (1.3%) to very common in The Gambia (64.9%). Non-synonymous mutants of *kelch13* were uncommon, most with unknown relevance to artemisinin resistance and observed for the first time in Africa. However, mutants that are artemisinin resistance-associated elsewhere were detected in three infection samples from Ghana (574L, 561H, 469Y), and one in Cameroon (538V). This large-scale genomic surveillance of *P. falciparum* infections highlights the need for ongoing monitoring of drug resistance and for data integration throughout the region.

## Introduction

Malaria caused by *Plasmodium falciparum* remains a major public health problem in West Africa, which accounts for nearly half of the global malaria burden, with an estimated 124 million cases in 2023 (1). Over the past two decades, significant progress has been achieved in reducing malaria morbidity and mortality, driven by effective treatment and chemoprevention strategies. Following the rise of resistance to previously used antimalarial drugs, artemisinin-based combination therapies (ACTs) were adopted as first-line treatments for uncomplicated malaria, and injectable derivatives of artemisinin or quinine for severe malaria (2). In 2012, the World Health Organisation (WHO) endorsed malaria chemoprevention for children under five at the start of transmission seasons in areas with seasonal transmission, with at least four treatment courses of sulfadoxine-pyrimethamine plus amodiaquine (SP +AQ). Since then, seasonal malaria chemoprevention (SMC) programs have been implemented across West Africa and have led to substantial reductions in malaria-related morbidity and mortality (3,4). Malaria chemoprevention for pregnant women and infants during periods of heightened vulnerability also relies on sulfadoxine-pyrimethamine (SP). More recently, rectal artesunate is recommended as pre-referral treatment of suspected severe malaria among young children (5). The efficacy of these treatment strategies is essential for sustaining malaria control and moving toward pre-elimination in West Africa.

Although current treatment strategies remain effective in West Africa, *de novo* emergence of ART-R has been documented in several countries across East and Northeast Africa (6,7). Similar threats also exist for chemoprevention strategies. Parasites circulating in West Africa already carry multiple resistance mutations to SP (8), and this background could enable them to develop resistance to amodiaquine component of the chemoprevention strategy. Early detection of emerging resistance mutations is therefore critical, as timely information on the rise of drug resistance can guide national malaria control programs (NMCPs) in implementing effective countermeasures.

Molecular surveillance of established genetic markers provides a systematic and scalable approach for monitoring antimalarial resistance (9,10). To support this effort, MalariaGEN developed an amplicon sequencing toolkit under the SpotMalaria project for genomic surveillance of malaria parasites using dried blood spots (DBS). This has been successful at genotyping samples from 24 countries (10–15). Briefly, the toolkit profiles each sample for a selected set of genetic markers, including genotypes associated with drug resistance, and a genetic barcode constituted from 100 single-nucleotide polymorphisms (SNPs) informative of parasite population structure and diversity. Implementation of the toolkit in regional sequencing hubs can significantly add value by integrating the data from satellite sites with standardisations in data processes (10,16).

Here, we present a new amplicon sequencing dataset comprising 17,384 *P. falciparum* samples from West Africa, sequenced at three independent sequencing centres (based in The Gambia, Ghana and the United Kingdom). We provide a comprehensive summary of this dataset by presenting: (i) prevalence of drug resistance markers with spatio-temporal patterns, (ii) low-frequency *kelch13* mutations of unknown importance, and (iii) parasite population structure. The genomic surveillance data per sample are presented in a standardised format known as Genetic Report Card (GRC)(10), enabling cross-sequencing-centre comparisons and facilitating future epidemiological analyses.

## Results

### Dataset summary

The West Africa GRC includes 17,384 *P. falciparum* infection samples collected from 57 first-level administrative divisions in eight countries. These samples were collected from 20 MalariaGEN studies (Table 1). The dataset exhibits wide heterogeneity in sample sizes, collection years, and geographic coverage across countries (Figure 1). Most samples (94.7%; 16,465/17,384) were collected between 2018 and 2023. Six of the eight countries represented in the dataset—except Guinea and Sierra Leone— contributed more than 30 samples collected in at least three different years. Over 80% of these samples showed longitudinal geographic overlap at the first- and second-level administrative divisions, except in Nigeria where only 32% and 9% of the samples showed overlap at the first and second administrative levels, respectively. Together, these characteristics provide a basis for the spatio-temporal analysis.

**Figure 1.**
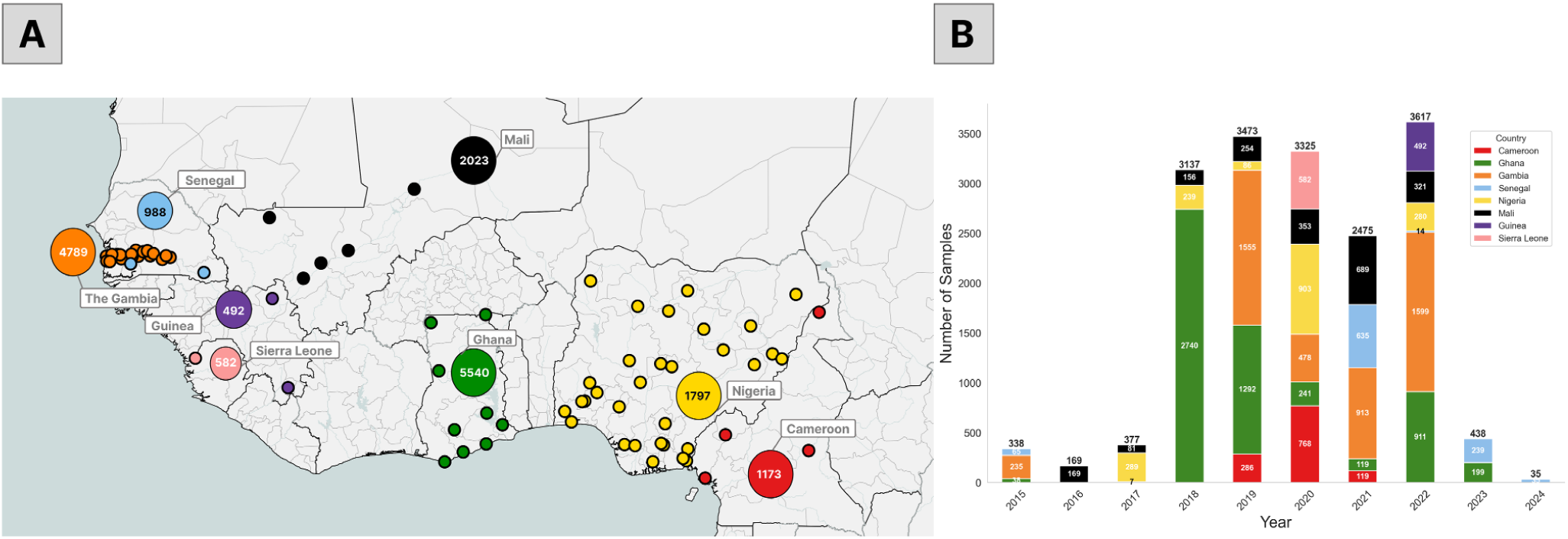
Geographic distribution of samples. **A.** Geographic distribution of samples collected from eight countries. Larger markers represent the sample counts per country, while smaller markers represent sampling sites at the second-level administrative division. **B.** Distribution of samples across countries in each year, colors correspond to the country color codes used in panel A.

**Table 1.** Summary of participating studies. Detailed descriptions of each study, including a key contact person, are provided at https://www.malariagen.net/project/spotmalaria/ and https://www.malariagen.net/project/genomic-surveillance-hubs-west-africa-nihr-global-health-research-group/. Sequencing of each sample was performed at one of three independent sequencing centres. Sample collection sites refers to the number of first and second-level (second-level shown in parentheses) administrative divisions that samples were collected by the respective study. Abbreviations: WACCBIP - West African Centre for Cellular Biology of Infectious Pathogens, University of Ghana, Ghana; MRCG - Medical Research Institute Gambia, London School of Tropical Hygiene and Medicine, The Gambia; WSI - Wellcome Sanger Institute, United Kingdom.

| MalariaGEN study ID | Sequencing centre | Country(s) surveyed | Sample collection sites | Sample collection years | Sample size |
| --- | --- | --- | --- | --- | --- |
| 1266 | WACCBIP | Ghana | 15 (30) | 2019 – 2020 –<br>2021 – 2022 -<br>2023 | 2476 |
| 1267 | MRCG | The Gambia | 8 (19) | 2017 – 2018 –<br>2019 – 2021 -<br>2022 | 2417 |
| 1374 |  | Senegal | 1 (1) | 2023 – 2024 | 274 |
| 1331 | WSI | Guinea | 2 (2) | 2022 | 492 |
| 1136 |  | The Gambia | 6 (8) | 2019 – 2020 | 1754 |
| 1332 |  | Sierra Leone | 1 (1) | 2020 | 582 |
| 1162 |  | The Gambia,<br>Senegal | 6 (6) | 2015 - 2022 | 555 |
| 1165 |  | Cameroon | 4 (4) | 2019 – 2020 -<br>2021 | 1173 |
| 1168 |  | Ghana | 1 (2) | 2018 | 2627 |
| 1182 |  | The Gambia | 1 (1) | 2015 | 135 |
| 1183 |  | Nigeria | 1 (1) | 2017 | 223 |
| 1197 |  | Mali | 3 (4) | 2016 – 2017 –<br>2018 – 2019 –<br>2020 -2021 –<br>2022 | 1535 |
| 1200 |  | Ghana | 1 (1) | 2015 – 2018 –<br>2019 | 212 |
| 1226 |  | Nigeria | 1 (1) | 2018 -2019 | 237 |
| 1294 |  | Ghana | 1 (1) | 2019 | 225 |
| 1306 |  | Nigeria | 24 (27) | 2017 – 2018 –<br>2019 – 2020 -<br>2022 | 1330 |
| 1316 |  | Mali | 1 (1) | 2021-2022 | 72 |
| 1317 |  | Mali | 2 (2) | 2021-2022 | 416 |
| 1318 |  | Senegal | 1 (1) | 2021-2022 | 376 |
| 1319 |  | Senegal | 1 (1) | 2021-2022 | 273 |

The complexity of infection (COI) index was estimated for 12,393 samples (71.3% of total) with at least 20 valid SNPs in their genetic barcode. The proportion of polyclonal samples, defined as COI>1, was determined for each country (Supplementary Table 1). The overall proportion of polyclonal infections in the region was 32.9%, ranging widely among different countries from 15.8% (The Gambia) to 57.9% (Cameroon). Notably, COI estimates based on THE REAL McCOIL algorithm are considered conservative (17), and therefore the proportion of polyclonal infections in West Africa is likely higher.

Mitochondria-based species detection identified non-falciparum *Plasmodium* species in 147 samples (0.84%), either detected as single-species (61.9%) or mixed-species (38.1%) infections. The other species detected in these samples were: *P. malariae* (97 samples; 0.56%) *P. ovale* (43 samples; 0.25%)*, P. vivax* (19 samples; 0.11%). Country-level distribution of detected species is summarised in Supplementary Table 2.

### *Plasmodium falciparum* population structure in West Africa

High-quality genetic barcodes of 5,879 samples were used to assess the parasite diversity and population structure in West Africa. Analysis of the 100-SNP genetic barcode revealed a regional average minor allele frequency (MAF) of 0.27 (Supplementary Figure 3), indicating that the SNPs within the genetic barcode are able to capture the allelic diversity in West African parasites. Pairwise genetic distance estimates of these samples were used for principal coordinate analysis (PCoA) to assess the population structure within the region. Parasites from different countries were not separated by the first two principal coordinates (Supplementary Figure 4A), suggesting substantial genetic mixing across the region. However, three distinct clusters of parasite populations from The Gambia were detected (Supplementary Figure 4A), with sample sizes of 91, 57, and 66. These clusters disappeared after repeating the PCoA on sample-pairs with pairwise genetic distance >0.1, suggesting that these clusters were formed by near-clonal parasite populations.

To assess the differentiation of the West Africa *P. falciparum* parasite population from other regions, PCoA was also performed on a combined dataset that included 5,368 samples from the Greater Mekong Subregion (GMS)—covering seven Southeast Asian countries—sourced from the publicly available GenRe-Mekong dataset (10). This analysis revealed a clear separation between West African and GMS samples along the first principal coordinate (Supplementary Figure 4B). This shows a potential future application of such genetic barcodes to detect any rare case of parasite importation between these two regions.

### Antifolate-resistance genotypic markers

The proportions of infections with *dhfr* triple-mutant alleles (51I/59R/108N)—used for determining resistance to sulfadoxine-pyrimethamine treatment (SP)— consistently exceeded 64% across all eight countries (Figure 2A, Table 2), indicating regional near-fixation of these mutations. The overall proportion of infections with SP resistance-associated alleles was 86.4%. Combinations of *dhps* and *dhfr* mutations, which are associated with higher levels of SP resistance, were also examined (Figure 3). Quadruple mutants, combining the *dhfr* triple mutations with the *dhps* 437G mutation, were also present in over 60% of samples across all eight countries. Samples harbouring the *dhfr* triple mutations and *dhps* double mutations 437G/540E were detected at low prevalence levels, ranging from 0% to 6.37% across countries over the years, with a regional prevalence of 1.82%. On the other hand, parasites carrying the combination of *dhps* 436A/437G and *dhfr* triple mutations were observed at a prevalence as high as 40% in Ghana, Nigeria, and Cameroon (Figure 3), with a regional prevalence of 29.3%. While the role of the 436A mutation in sulfadoxine resistance is not fully established, its presence alongside other mutations may contribute to reduced drug efficacy (17).

**Figure 2.**
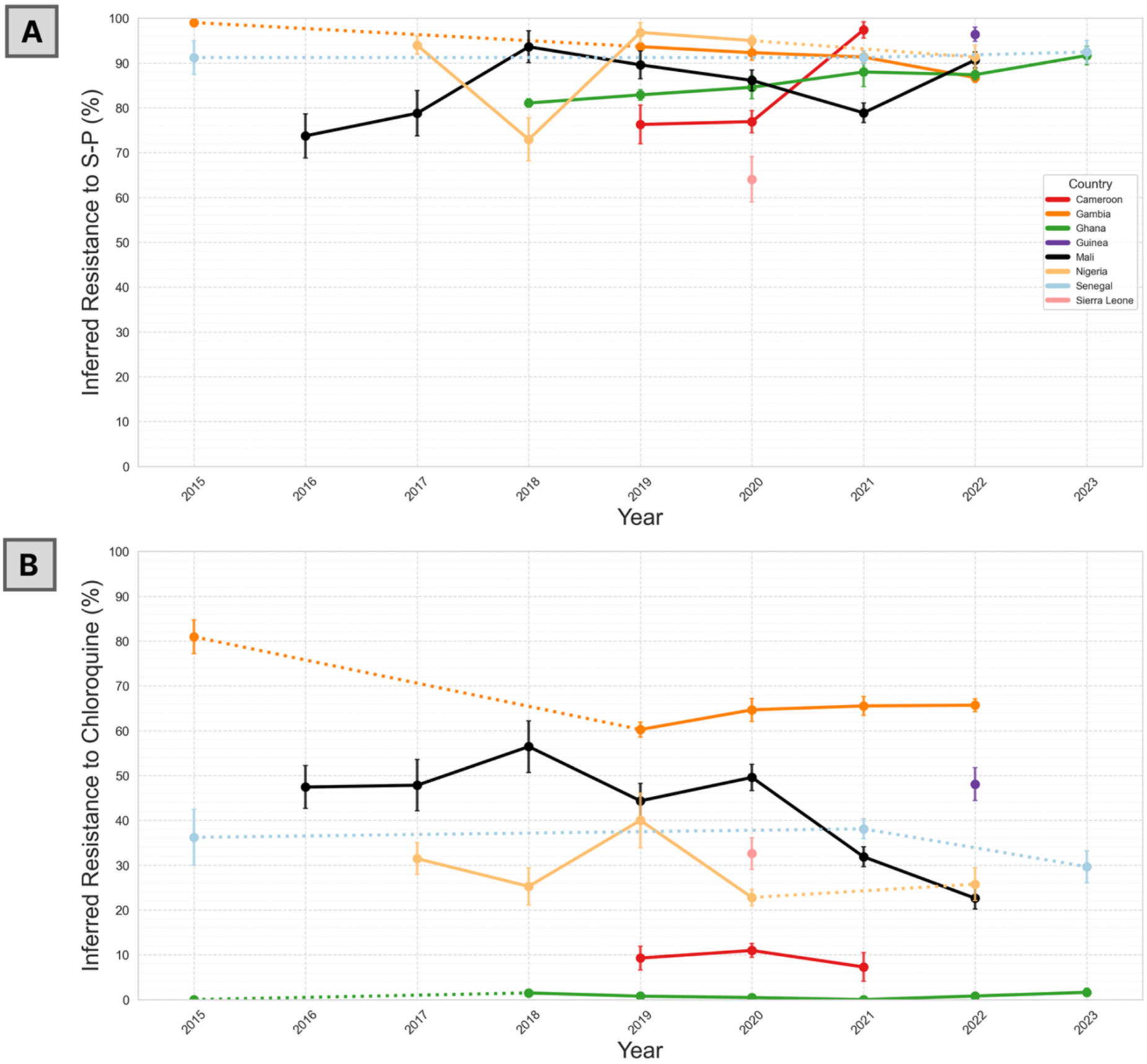
Prevalence of inferred resistant samples across countries over time. Vertical error bars indicate the standard error of inferred resistance prevalences for a given year. Only countries with at least 30 samples are shown. A. Inferred resistance to sulfadoxine-pyrimethamine, defined by the presence of *dhfr* N51I/C59R/S108N mutations. B. Inferred resistance to chloroquine, defined by the presence of the *crt* K76T mutation.

**Figure 3.**
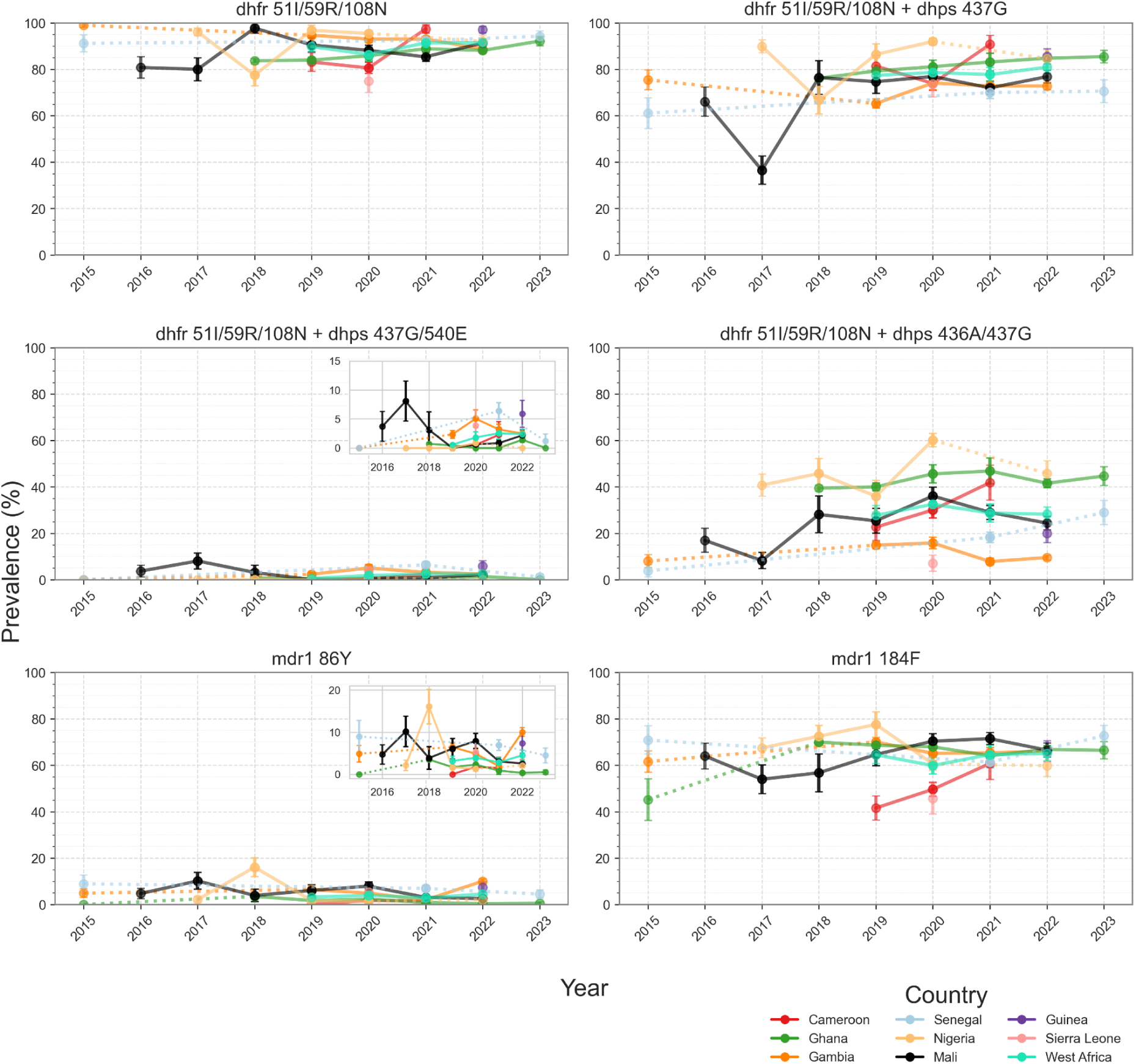
Temporal changes in the prevalence of *dhfr/dhps* combined mutations and *mdr1* N86Y, Y184F mutations in West African countries. Vertical error bars indicate the standard error of inferred resistance prevalences for a given year. Samples with missing or heterozygous calls in the respective haplotypes were excluded from the analysis. Prevalence was calculated regardless of other mutations in the respective gene. Only countries with at least 30 samples at two or more time points are shown in the plot.

**Table 2.** Drug resistance-related mutations genotyped. The details of the amplicon primers for each genotyped loci is provided in the Supplementary File 2. *Mutations used to predict resistance phenotype to the corresponding antimalarials.

| Gene Description | Mutation | Antimalarial |
| --- | --- | --- |
| *kelch13* | any mutation seen in BTB/POZ and propeller domains, including WHO list of validated/candidate mutations* | Artemisinin |
| *dhfr*<br>(bifunctional dihydrofolate reductase-thymidylate synthase) | N51I* | Pyrimethamine |
|  | C59R*/Y |  |
|  | S108N*/T |  |
|  | I164L |  |
| *mdr1*<br>(multidrug resistance protein 1) | N86Y | Chloroquine, Amodiaquine, Lumefantrine, Mefloquine |
|  | Y184F |  |
|  | D1246Y |  |
| *crt*<br>(chloroquine resistance transporter) | C72S | Chloroquine |
|  | M74I |  |
|  | N75D/E |  |
|  | K76T* |  |
|  | N326S | ART-R genetic background |
|  | I356T |  |
| *dhps*<br>(dihydropteroate synthetase) | S436A/Y/F/G | Sulfadoxine |
|  | A437G* |  |
|  | K540E/N |  |
|  | A581G |  |
|  | A613S/T |  |
| *exo* (exonuclease) | E415G | Piperaquine |
| *pm23* (plasmepsin 2 and plasmepsin 3) | Breakpoint |  |
| *fd* (ferredoxin) | D193Y | ART-R genetic background |
| *mdr2*<br>(multidrug resistance protein 2) | T484I |  |
| *arps10*<br>(apicoplast ribosomal protein S10) | V127M |  |
|  | D128Y/H |  |

None of the parasites carried the *dhps* 437G/540E/581G or the *dhps* 437G/540E/613S triple mutations. However, *dhps* 436A/437G/581G/613S mutations were present across the countries, with a regional prevalence of 7.6%. These *dhps* mutations were notably more prevalent in Nigeria (29.7%) and Cameroon (12.2%) than in the other West African countries (2.0%).

### Chloroquine and amodiaquine inferred resistance markers

The regional prevalence of chloroquine resistance, inferred from the presence of *crt* 76T mutation, was 25.5%. It was greater than 10% across all countries except Ghana where resistance is considerably lower (1.1%) compared to the rest of the region. The prevalence of chloroquine resistance (64.7%) in The Gambia was the highest in the region (Table 3). However, the prevalence fluctuated over the years, decreasing from 80.9% in 2015 to 60.2% in 2019 (adjusted p-value [adj-p] <0.001, odds ratio [OR]=2.8), and then rising to 64.6% (adj-p=0.6, OR=0.8) in 2020 (Figure 2B). A contrasting pattern was observed in Mali, where the prevalence of chloroquine resistance initially increased from 47.8% in 2017 to 56.4% in 2018 (adj-p=0.7, OR=0.7), before declining to 31.8% in 2021 (adj-p <0.001, OR=2.1) and further declining to 22.6% in 2022 (adj-p =0.08, OR=1.6).

**Table 3.**
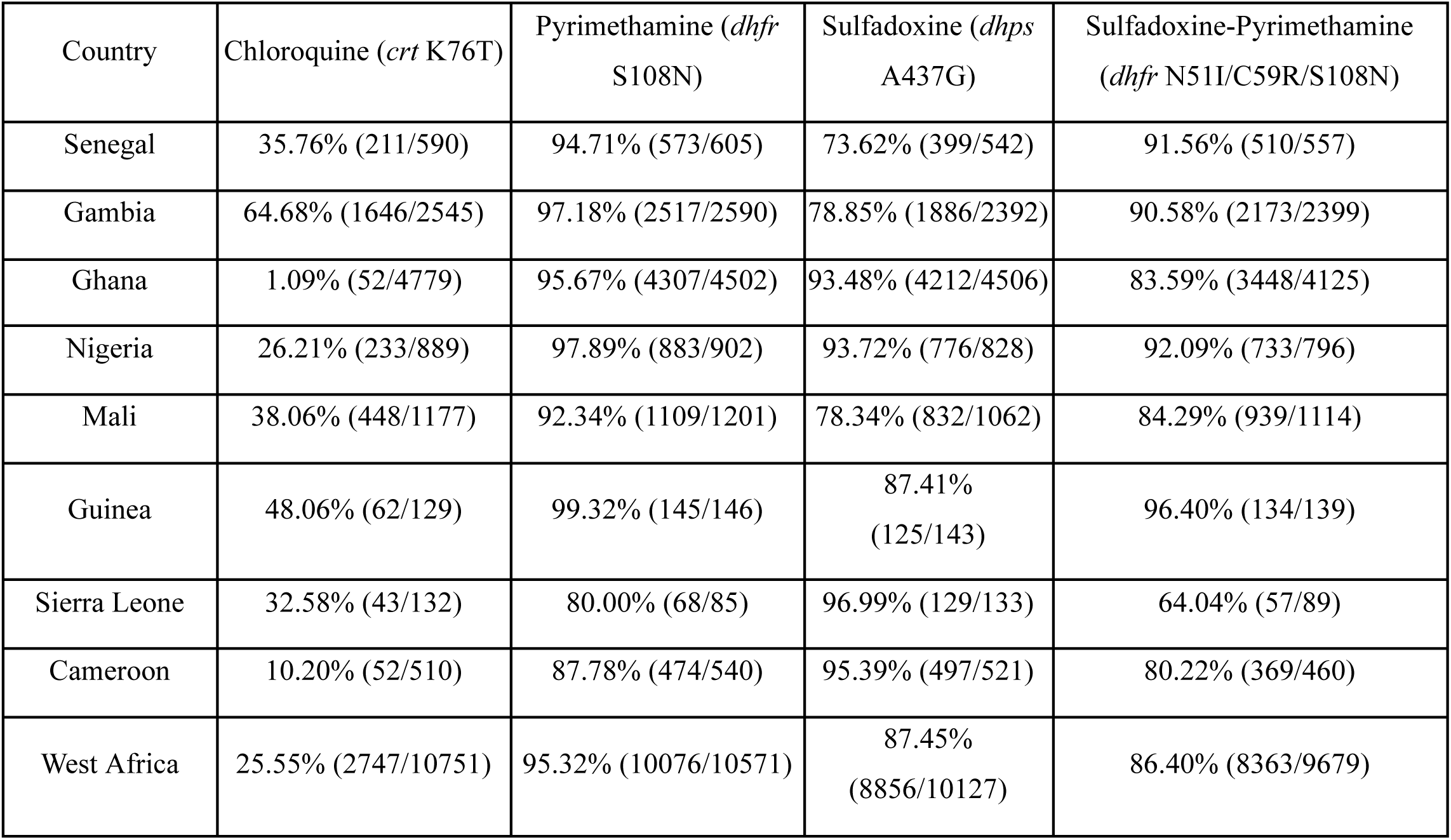
Prevalence of drug resistance across countries. Sample counts are included in the parenthesis. Mutations used for phenotype predictions were listed in Table 2. Artemisinin and Piperaquine are excluded from the table since the prevalence of drug resistance was lower than 1% for both drugs across the countries.

| Country | Chloroquine (*crt* K76T) | Pyrimethamine (*dhfr* S108N) | Sulfadoxine (*dhps* A437G) | Sulfadoxine-Pyrimethamine (*dhfr* N51I/C59R/S108N) |
| --- | --- | --- | --- | --- |
| Senegal | 35.76% (211/590) | 94.71% (573/605) | 73.62% (399/542) | 91.56% (510/557) |
| Gambia | 64.68% (1646/2545) | 97.18% (2517/2590) | 78.85% (1886/2392) | 90.58% (2173/2399) |
| Ghana | 1.09% (52/4779) | 95.67% (4307/4502) | 93.48% (4212/4506) | 83.59% (3448/4125) |
| Nigeria | 26.21% (233/889) | 97.89% (883/902) | 93.72% (776/828) | 92.09% (733/796) |
| Mali | 38.06% (448/1177) | 92.34% (1109/1201) | 78.34% (832/1062) | 84.29% (939/1114) |
| Guinea | 48.06% (62/129) | 99.32% (145/146) | 87.41% (125/143) | 96.40% (134/139) |
| Sierra Leone | 32.58% (43/132) | 80.00% (68/85) | 96.99% (129/133) | 64.04% (57/89) |
| Cameroon | 10.20% (52/510) | 87.78% (474/540) | 95.39% (497/521) | 80.22% (369/460) |
| West Africa | 25.55% (2747/10751) | 95.32% (10076/10571) | 87.45% (8856/10127) | 86.40% (8363/9679) |

The *crt* 74I/75E/76T (CVIET) haplotype, when combined with *mdr1* 86Y, has been associated with resistance to amodiaquine in West Africa (18,19). The vast majority of samples with a homozygous haplotype for codons 72-76 had either CVIET (regional prevalence of 31.1%) or the wild-type CVMNK (68.8%). As such, the prevalence of CVIET closely matches that of 76T alone. The SVMNT haplotype, which mediates amodiaquine resistance in Asia and the Americas (20), was not detected.

### *mdr1* alleles affecting partner drug susceptibility

Mutations in *mdr1* at codons 86Y, 184F and 1246Y have been associated with altered susceptibility to lumefantrine and amodiaquine in sub-Saharan Africa (21,22). Specifically, the wild-type N86 (95.8%) has been associated with lumefantrine resistance, while the mutant 86Y (4.2%) has been associated with amodiaquine resistance. The regional prevalence of other *mdr1* mutations 184F and 1246Y were 63.5% and 0.42%, respectively. Notably, *mdr1* N86/184F/D1246 genotype was present in 59.7% of the parasites, likely reflecting the selection pressure exerted by AL treatment in the region. Samples carrying triple *mdr1* mutations 86Y/184F/1246Y were not observed in the region. Country-level temporal assessments of these mutations are shown in Figure 3. In The Gambia, the prevalence of 86Y increased from 1.9% in 2021 to 10% in 2022 (adj-p<0.001, OR=0.18), warranting continued monitoring of this mutation in the following years. Another significant increase in the prevalence of the 86Y mutation was seen in Nigeria, rising from 1.5% in 2017 to 16.0% in 2018 (adj-p=0.002, OR=0.11). However, this increase was sporadic and likely due to sampling heterogeneity, as the prevalence dropped back to 1.7% (adj-p=0.02, OR=11.3) and 1.3% (adj-p=0.8, OR=0.6) in the following years. *mdr1* 184F mutation was observed at over 50% prevalence across countries for most years. A continuous increase in the prevalence of *mdr1* 184F in Cameroon between 2019 and 2021 was noted (adj-p>0.05, OR=0.6, Figure 3).

### Detection of potential artemisinin resistance-associated *kelch13* mutants

Among 9,079 (52.7%) *P. falciparum* samples successfully genotyped for the BTB/POZ and propeller regions of *kelch13*, 168 samples carried a non-synonymous codon change (1.8%), observed either as heterozygous (90 samples; 1.0%) or homozygous (78 samples; 0.9%). Mutations observed were annotated as to whether they were reported in the global dataset of *kelch13* mutations described by Balmer et al. (2025) (23) and in the WHO compendium of molecular markers for antimalarial drug resistance (24). There were 31 unique mutations in *kelch13* (Figure 4), 15 were detected for the first time in Africa, and 12 were detected for the first time globally. 17 of the 31 mutations were seen in only a single sample. The most common mutation was 578S, which is not considered to be associated with ART-R.

**Figure 4.**
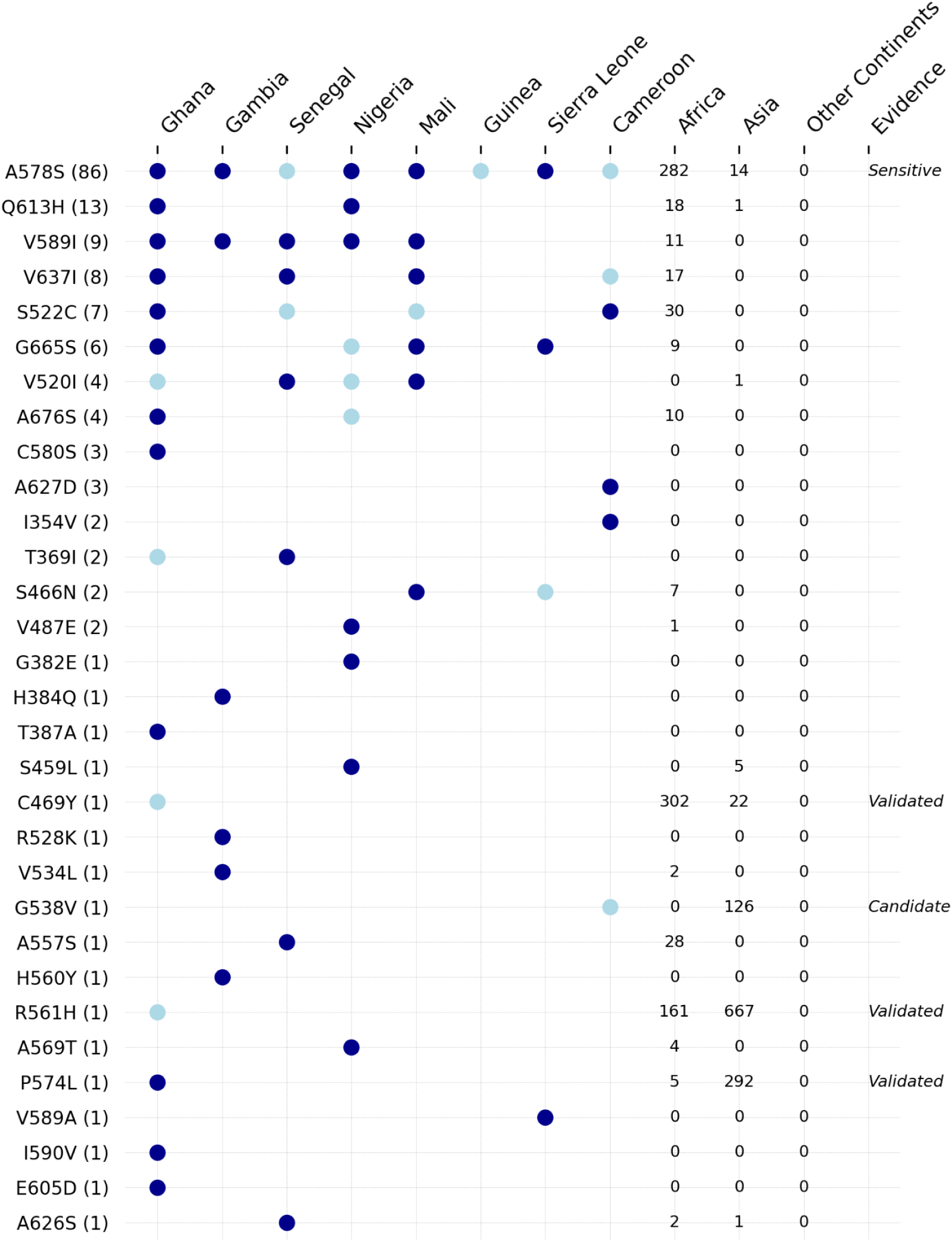
Country breakdown of 31 non-synonymous *kelch13* mutations observed in the dataset. Cumulative count of each mutation (y-axis) is denoted within the parentheses, and count of the same mutation in the global *kelch13* mutations dataset (Balmer *et al*., 2025) was also reported in the latter columns (namely Africa, Asia and Other continents). Samples from this study were not counted in the Africa column. Dark-blue points denote homozygous calls; light-blue points denote heterozygous calls. If a mutation is observed as both homozygous and heterozygous in a country, it is rendered dark blue. Heterozygous calls are shown only for mutations that appear at least once as homozygous in our dataset and/or WHO validated/candidate markers. The last column indicates the ART-R evidence status of a mutation according to the WHO.

Four ART-R-associated markers were detected, each occurring in a single sample: three from Ghana, and one from Cameroon. All three ART-R markers 469Y, 574L and 561H that were observed in Ghanaian samples were previously reported in East Africa (25–27). On the other hand, the ART-R-associated marker 538V found in a sample from Cameroon is reported here for the first time in Africa. Of these four mutations, only 574L was seen as a homozygous call, with the other three mutations occurring together with wild-type allele.

### Assessment of genetic backgrounds supporting *kelch13* mutations

The co-existence of *kelch13* mutations with mutations in other drug resistance genes (*crt, mdr1, dhfr* and *dhps)* was examined to assess whether a specific genetic background might support the emergence of *kelch13* mutations. Comparisons were conducted at the country-level using Fisher’s exact test to evaluate differences between parasites carrying wild-type versus *kelch13* mutants. No statistically significant (<0.05 after Benjamini-Hochberg correction) differences were observed between the two groups in any country (Supplementary Figure 1).

The mutations in *mdr2, crt, ferredoxin, arps10* that were observed in the genetic background of *kelch13* mutations in Asia (28) were also surveyed (Table 2). The *crt* 356T mutation, previously associated with the artemisinin resistance genetic background in Southeast Asia (28) and quinine resistance in West Africa (29), was detected in 2,107 samples (regional prevalence of 24.6%). Its prevalence was greater than 30% in countries across the western part of the region—Gambia, Senegal, Mali, and Sierra Leone— and low in Ghana (0.5%), and at intermediate frequencies in Nigeria (20.4%) and Cameroon (6.9%), largely reflecting frequencies of 76T (Supplementary Figure 2). It might be important to determine whether this mutation could play a role in the background of *kelch13* mutations in West Africa. Other known “background” mutations were either absent or occurred at very low frequencies (<1%).

## Discussion

Effective treatments of malaria and targeted interventions such as chemoprevention programmes put parasite populations under heavy drug selection pressure. Genomic surveillance can detect adaptations towards drug resistance and inform decision-making of national malaria control programs (10). Here, we present the largest regional landscape analysis of antimalarial resistance in West Africa to date. The collaboration within the subregion has enabled the processing of 17,384 *P. falciparum* samples collected between 2015 and 2024 across eight countries. This number is 4,000 more than previous data from the largest whole genome sequences of 13,000 genomes from West Africa in the MalariaGEN Pf8 whole-genome data resource (30). This highlights rapid enhancement of data facilitated by close collaborative effort between scientists in the malaria surveillance ecosystem with national malaria control programmes. The data also go beyond national boundaries to harness the power of data integration at regional scales to understand the broader architecture of circulating drug resistance variants, correlation with front-line therapies, and other targeted interventions such as SMC, PMC etc for broader strategic regional policy direction. This is against the backdrop of efforts being made by the AfricaCDC Pathogen Genomics Initiative (PGI) to harmonize and federate surveillance products and priorities within the ecosystem to ease data integration at a greater continental scale. In West Africa, the use cases are imminent as the region keeps watch against any emergence of artemisinin partial resistance, but also in monitoring current important mass drug interventions such as seasonal malaria chemoprevention (SMC) with SP + amodiaquine or the use of SP alone in IPTp.

As SP-based chemoprevention expands in the region (31), continuous drug pressure will likely increase the risk of persistence of legacy resistance to these drugs, with the risk of evolution of new multi-drug resistant variants. Currently, *dhfr/dhps* quadruple mutants are circulating at high prevalence (76%) in West Africa, but SP remains largely efficacious in chemoprevention (32), given the low prevalence of the common *dhps* A437G/K540E/A581G mutations that confer a more resistant phenotype in East Africa. Another *dhps* haplotype that carries a greater risk of resistance is the 431V/436A/437G/581G/613S quintuple mutation, first documented in 2007 in Nigeria (33), later reported in Mali (34), Burkina Faso (35), Cameroon (36) and elsewhere in Central Africa (37). In our study, the prevalence of 436A/437G/581G/613S mutations was notably higher in Nigeria (29%) and Cameroon (12%) than in the rest of West Africa (2%). The current version of the amplicon panel does not genotype the 431V mutation, but analysis of the MalariaGEN Pf8 whole-genome data resource using Pf-HaploAtlas reveals that this mutation is linked to the quintuple haplotype in 97.4% (149/153) of the samples with the haplotype (38). Although the exact impact of these mutations are unknown, evidence associating them to resistance to sulfadoxine is increasing (35,37,39). Future studies assessing their effect on the efficacy of SP in West Africa will be valuable in planning future chemoprevention strategies.

The two most commonly used ACTs in Africa, artemether–lumefantrine (AL) and artesunate–amodiaquine (AS-AQ), exert opposing selection pressures on the *mdr1* gene. In particular, the presence of *mdr1* N86 and/or D1246 genotypes results in increased tolerance to lumefantrine, while *mdr1* 86Y and/or 1246Y genotypes increase tolerance to amodiaquine (18,40,41). In this dataset, the *mdr1* mutations N86Y (4.2%) and 1246Y (0.5%) were low in prevalence, and the wild variants are selected by lumefantrine, suggesting possible increased tolerance to lumefantrine in circulating parasites. Lumefantrine resistance has recently been associated with variants in the *px1* gene (PF3D7_0911500) (42), but the frequency of these variants are rare in West Africa, an indication that alternative pathways to lumefantrine resistance may be operating in the region. The *mdr1* 184F has been increasing in frequency in this region, and is known to be associated with lumefantrine tolerance (41). On the other hand, half of the parasites carrying the 86Y mutation had *crt* CVIET haplotype, known to have increased tolerance to amodiaquine (18). These together support the continuous use of amodiaquine in first-line ACTs and SMC, but warrant enhanced monitoring and experimental evaluation of the impact of evolving *mdr1* and emerging *px1* variants on artemether-lumefantrine efficacy across West African populations.

Following the official discontinuation of chloroquine as a first-line treatment for uncomplicated malaria in West Africa, a declining trend was reported for *crt* 76T across most endemic countries (30). However, the prevalence of chloroquine-resistant parasites in West Africa remains greater than 10% across countries, except Ghana, where the vast majority of parasites carry the 76K wild variant of *crt* and thus inferred to be now susceptible again to chloroquine. In the absence of chloroquine pressure, other factors may be sustaining pressure on *crt,* such as the use of amodiaquine as part of SMC programs (43). Exceptionally, the *crt* 76T mutation persists at the highest frequency in The Gambia (44–46), where parasite prevalence is now very low, with clonal expansion and divergent founder sub-populations observed. This high persistence is common in other far-west African countries like Senegal (30), probably due to compensatory variants in these populations, that are yet to be determined. Although ACTs remain clinically efficacious in the region, the impact of this persistent chloroquine resistance on chemo-interventions or the risk to the efficacy of ACTs and candidate drugs needs further investigation.

The emergence of ART-R in Africa is a significant concern to malaria control and elimination. While ART-R markers have been increasingly reported in East Africa, observations in Central and West Africa have been sporadic (23). Similarly, we identified four ART-R associated markers (574L, 561H and 469Y in Ghana and 538V in Cameroon), each observed in a single sample. While only 574L was observed in a homozygous infection (318 reads), other markers also had strong evidence (>15 reads, >50% of total alleles). Overall, 1.8% (168/9,079) of the parasites surveyed carried 31 non-synonymous mutations in *kelch13*, mostly the 578S mutant not associated with ART-S, or other mutants of unknown clinical importance and many observed for the first time in Africa. Although these variants of unknown clinical importance are generally assumed to be mostly deleterious and often disappear, the variety detected in the region presents a risk to independent ART-R emergence in West Africa, as has been seen in East Africa (47). The emergence of these sporadic mutations in *kelch13* highlights the need for ongoing surveillance of drug resistance and further research into their effects on the biology, treatment and transmission of *P. falciparum* in the region.

Understanding why the prevalence of drug resistance varies across countries in the region is crucial for regional assessments. One key factor could be the differential selection pressure exerted by various antimalarial drugs, as different medications favour the emergence and selection of different resistant parasite variants. The adoption and availability of different artemisinin-based combination therapies (ACTs) play a significant role in this dynamic. Additionally, differences in the timing of first-line treatment policies, the introduction of either multiple or single ACTs, and the implementation of seasonal malaria chemoprevention (SMC) programs—often rolled out in different years—may contribute to these disparities (48). Variations in malaria transmission intensity in West Africa, driven by ecological, vector species and other bioclimatic conditions, are likely to have further influenced the patterns of drug resistance observed. In areas of high and seasonal transmission, a large proportion of infections circulate among asymptomatic or mildly symptomatic individuals who do not seek care due to higher levels of immunity or geographical, financial and structural barriers, which could reduce the overall drug pressure (49). Meanwhile, the push for elimination in low transmission settings like The Gambia and Senegal, allow for higher drug pressure, e.g. from SP-amodiaquine use in SMC, which could be maintaining the legacy mutations. Also, human and vector genetic variations that influence within-host dynamics, including variable drug metabolism in humans, may be playing a role (50).

There are limitations in this study related to the data and the analysis methods. The data were generated from samples collected in independent studies with varying designs and uneven spatio-temporal sampling (Figure 1). This resulted in uneven geotemporal representation of parasite populations across countries. Although one of the advantages of genomic surveillance is the high throughput in covering a large number of samples for drug resistance inference, more information on the clinical relevance of the patterns observed will be drawn from accompanying this with clinical and laboratory studies of drug susceptibility. Moreover, the current survey only targets known genetic markers of resistance, while resistance to antimalarials may arise through alternative other mechanisms (51), sometimes driven by the background of the parasite population)(52). Therefore, expanding to genome-wide surveillance and genotype–phenotype association studies as malaria parasites continue to evolve against interventions and the environment, may provide more accurate assessments of local drug resistance status.

Moving forward, continuous and expanded collaborations between the scientists and National Malaria Control Programs (NMPCs), incorporating assessment of parasites post drug interventions or efficacy tests are essential. Such coordinated efforts could maximise existing sequencing capacity, harmonise laboratory and data reporting standards, streamline sampling strategies and operational processes, and integrate genomic surveillance with routine malaria surveillance platforms, towards early detection of antimalaria drug resistance threats in the population in West Africa. The data generated in this study are suitable for integration into the wider data platforms serving as a catalogue for malaria drug resistance research (53,54). In summary, the landscape of *P. falciparum* drug resistance in West Africa presented by this study underscores the need for continuous genomic surveillance across Africa and offers insights for malaria control strategies.

## Methods

### MalariaGEN partner studies

Samples were obtained from 20 independent MalariaGEN studies (Table 1) in seven West African countries (The Gambia, Senegal, Sierra Leone, Guinea, Mali, Ghana, Nigeria), as well as Cameroon, a neighboring country in Central Africa sharing a large border with Nigeria. Although Cameroon is geopolitically located in Central Africa, it was included in this dataset because parasite populations from this area are genetically similar to those circulating in West Africa, as shown in previous large-scale analyses of *P. falciparum* population structure (30). Details of these studies are available at https://www.malariagen.net/project/genomic-surveillance-hubs-west-africa-nihr-global-health-research-group/ and https://www.malariagen.net/project/p-falciparum-community-project/. As studies have differences in sampling strategies, a general overview of the sample collection procedure is provided here. Samples with the following MalariaGEN study IDs were previously published elsewhere: 1197 (55,56), 1306 (57), 1316-1317 (58), 1318-1319 (59).

### Sample collection

Participants recruited at healthcare facilities were initially screened for *Plasmodium falciparum* (*Pf*) malaria infection either with rapid diagnostic tests and/or microscopy. Following the consent of the individual or their parent/guardian, samples confirmed positive for *P. falciparum* were collected as dried blood spots (DBS) by finger-prick. Samples were labelled with an anonymous identifier and registered in a secure bespoke online platform along with minimal metadata for tracking purposes. The DBS samples were then shipped to one of the three sequencing hubs (see below for more details) for the downstream processing and genomic analysis. Ethical approvals were obtained from ethics committees in each country of collection (listed in Consent section).

### Sample processing

Samples were processed following the protocols designed for MalariaGEN Amplicon Toolkit, implemented in laboratories within the following institutions: West African Centre for Cellular Biology of Infectious Pathogens (WACCBIP), University of Ghana, Ghana; the Medical Research Institute Gambia (MRCG) at London School of Hygiene and Tropical Medicine, The Gambia; and the Wellcome Sanger Institute (WSI), United Kingdom.

Across all sites, DNA extraction was performed according to the manufacturer’s instructions using the Qiagen DNA Investigator Kit. Extracted parasite DNA was amplified by applying selective whole-genome amplification (sWGA), using the methodology previously described by Oyola et al. (60). Amplicons underwent a two-step PCR protocol, with PCR1 amplifying target amplifications in multiplex PCR reactions immediately followed by PCR2 where bespoke index tags are ligated which allows the clusters derived from different samples to be unambiguously identified on the MiSeq flowcell. The final libraries were designed to capture a narrow size range (190-250bp inclusive of primer sites) to complement the sequencing length of the MiSeq v2 300 kit (Illumina, San Diego, CA). Fragment sizes confirmed using the Agilent Tapestation system using Genomic DNA High Sensitivity D1000 Screentape. A second quality control step was used to quantify the library concentrations by qPCR reaction. Sample libraries were sequenced using the Illumina MiSeq platform using the 150 PE MiSeq run -Reagent Kit v2 (300 cycle). Manual laboratory workflows were implemented at the West African hubs for data generation, and high-throughput automated laboratory workflows were carried out at the WSI. Detailed protocols for the *P. falciparum* Amplicon manual laboratory workflow are available on the MalariaGEN Amplicon Toolkit Resources page: (https://www.malariagen.net/resources/amplicon-sequencing-toolkit/p-falciparum-amplicontoolkit-protocols/)

### Genotyping

Genotypes were called using the publicly available AmpRecon bioinformatics pipeline version 1.4.0 (https://github.com/malariagen/AmpRecon/). Details on the genotyping methods are provided in the *SpotMalaria Technical Notes and Methods* published as supplementary material in Jacob *et al*. (10), available from the “SPOTmalaria Technical Notes and Methods” link at https://www.malariagen.net/resource/29/. Briefly, genotype calls were based on read-count thresholds. A homozygous genotype required at least 10 high-quality reads supporting a single allele. A heterozygous genotype required at least 5 high-quality reads supporting the minor allele, with the minor allele representing a minimum of 10% of the total reads at that position.

A survey of drug resistance mutations was carried out within genes *mdr1*, *dhfr, crt, dhps, kelch13* (Table 2), and a further six mutations that were associated in the background of *kelch13* mutations in Southeast Asia (28). For *kelch13,* any non-synonymous mutation seen in BTB/POZ and propeller domains were genotyped. Subsequently, an extra curation step excluded non-synonymous heterozygous mutations in *kelch13* that have not been genotyped as homozygous in any of the samples analysed here.

100 SNPs that are informative of the *P. falciparum* parasite diversity are genotyped and concatenated to form a genetic barcode (the details of the SNPs are provided in the Supplementary File 2). Parasite mitochondrial-reads were used to confirm the infecting species. Samples identified as non-*P. falciparum* or mixed-species infections were excluded from the analysis set. However, because the sWGA approach used does not specifically target mitochondrial DNA, samples lacking mitochondrial reads—and therefore species calls—were still retained.

### Genetic report card

Sample metadata and genotype calls from individual multiplex runs were stored in a GRC. This is structured as a tabular file, where each row represents a particular sample. The following field groups are presented as columns: metadata fields, sample selection, drug resistance predictions, sample characterization, key drug resistance genotypes and haplotypes, detailed genotypes at drug resistance loci, genotypes used for genetic barcodes. The GRC provided with this study is generated by combining the GRCs generated for each participating study, and is provided in the Supplementary File 3.

### Methods for drug resistance and allele prevalence calculations

Sample drug resistance phenotypes were inferred using genotype-based rules previously described by Jacob *et al*. (see “Phenotype rules” link at https://www.malariagen.net/resource/29/), classifying each sample as “resistant”, “sensitive” or “undetermined” for the five antimalarial treatments: Chloroquine, Sulfadoxine, Pyrimethamine, Sulfadoxine-Pyrimethamine (SP), Artemisinin. Drug resistance and allele prevalence estimates were calculated by aggregating samples at the country level after excluding those with undetermined phenotypes or heterozygous calls at the relevant resistance loci.

All reported prevalence estimates were based on datasets containing a minimum of 30 samples. Sample aggregations for prevalence calculations were performed at three levels:

1. Country-level prevalences, based on samples aggregated from the respective country.
2. Country-year-level prevalences, based on samples aggregated by country-year pairs.
3. Regional-level prevalences, calculated as the simple unweighted mean of country-year prevalences across the eight countries.

Differences in drug resistance and allele prevalence between adjacent sampling years within each country was assessed using Fisher’s exact test. To account for multiple comparisons, p-values were adjusted using the Benjamini-Hochberg procedure separately for each genetic marker. Uncertainty for country-level and country-year level resistance prevalences were quantified using binomial standard errors, whereas for regional-level prevalences standard error of mean was reported.

### Methods for genetic barcode analysis

The complexity of infection (COI) of individual samples were inferred using the REAL McCOIL (61) in samples with less than 80% missingness in their barcode. For population structure analysis, samples with less than 10% missingness in their genetic barcode and a complexity of infection (COI) estimate equal to 1 were retained. Pairwise genetic distances between samples were calculated using the grcMalaria package (10), based on the barcode data, while excluding missing genotype calls. Samples with a pairwise genetic distance of less than 0.1 to the 3D7-derived genetic barcode were excluded to minimize the risk of contamination. Principal coordinate analysis (PCoA) based on the pairwise genetic distances was used to estimate the population structure within West Africa.

Visualisations were generated using a combination of tools. The grcMalaria R package was used to create maps, and other figures were created using customised Python (version 3.9) scripts.

### Data availability

The data used in this paper, including all sample metadata and genotypes, are openly available at Supplementary File 3. Amplicon raw sequence data were deposited in the European Nucleotide Archive (ENA) under accession numbers PRJEB85801, PRJEB2136, PRJEB94802.

### Consent

Ethical approval for the studies that contributed samples to this dataset was given by the following local and institutional committees:

Institutional Review Board of the Faculty of Health Sciences at the University of Buea Cameroon, Cameroon; Ghana Health Service Ethical Review Committee, Ghana; The Gambia Government/MRC Joint Ethics Committee, Banjul, The Gambia; Cross River State Health Research Ethics Committee, Cross River State Ministry of Health, Calabar, Nigeria; Federal Medical Centre, Yola, Nigeria; Comite d’Ethique de la FMOS/FAPH Universite des Sciences, des Techniques et des Technologies de Bamako and the Comité National pour la Recherche en Santé in Sénégal, Senegal.

### Funding

This research was partly funded by the NIHR (NIHR134717 and Kwiatkowski 17/63/91) using UK international development funding from the UK Government to support global health research. The views expressed in this publication are those of the author(s) and not necessarily those of the NIHR or the UK government. The Ghana sample collections were supported by the Gates Foundation (INV-050873). The Gambian collections were supported by the Pan-African Malaria Genetic Epidemiology Network, EGSAT and Genomic Surveillance Hubs in West Africa projects funded by the Wellcome Trust, EDCTP2 and NIHR, UK respectively.

## Acknowledgements

We would like to thank Frank Schwach, Andrea Frick-Kretchschmer and Bruhad Dave for their support with bioinformatics processes; Christen Smith and Rachel Wuendrich Ogidan for their assistance with compliance procedures; and Chiyun Lee, Nina White and Andrew Balmer for their feedback on visualisations. We are also deeply grateful to the patients who participated in these studies and the medical staff at these facilities for their invaluable support in patient screening.

