## Supplementary Figure for "Characterising recent antimalarial resistance in West Africa: Insights from amplicon sequencing of 17,384 *Plasmodium falciparum* infection samples"

### Supplementary Material

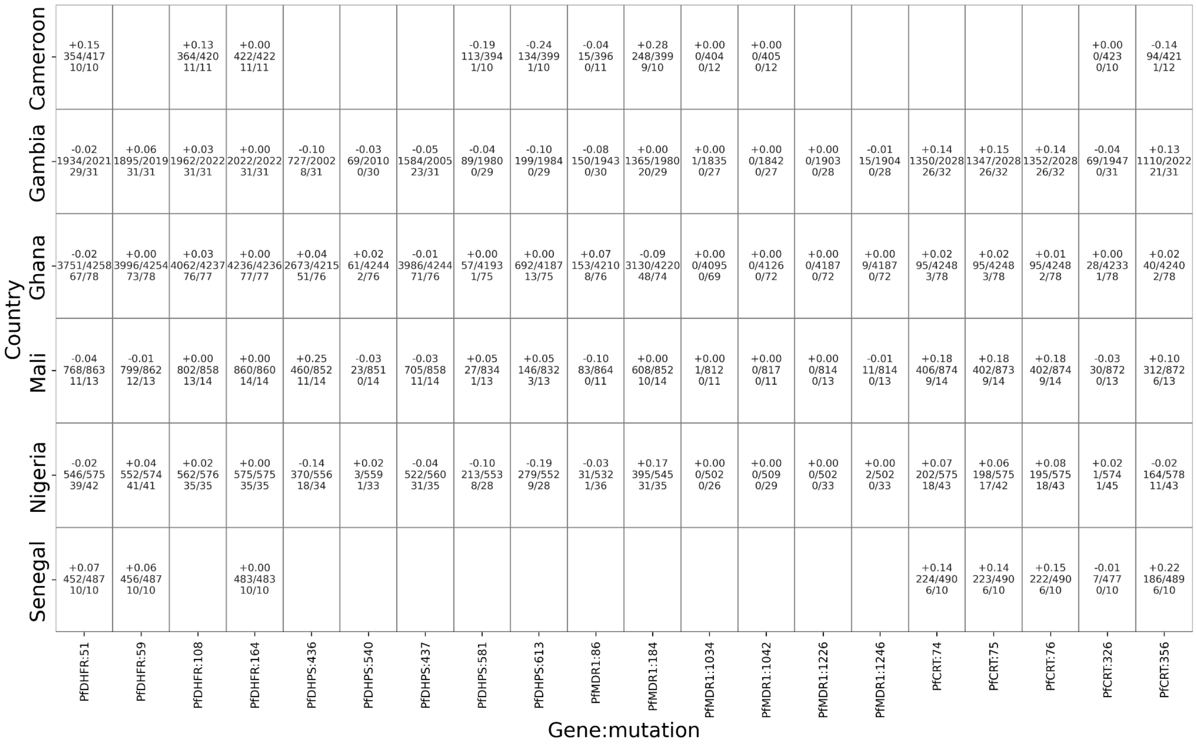


**Supplementary Figure 1. Comparison showing the prevalence of drug resistance-related mutations in samples carrying *kelch13* mutations (either as homozygous or heterozygous) and *kelch13* wild-type across countries.** Comparison is only shown if at least 10 samples were present in each group. Each box contains three rows: difference in prevalence (%) between *kelch13* mutant and wild-type groups, number of samples with the mutation in the *kelch13* wild-type group, number of samples with the mutation in the *kelch13* mutant group.

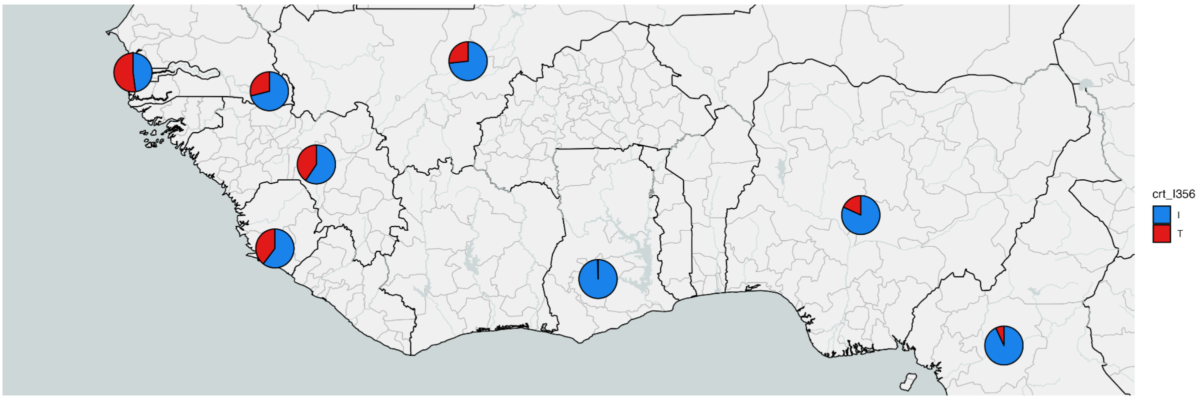


**Supplementary Figure 2. Presence of *crt:*356T allele in countries in West Africa.** Each pie chart refers to allele prevalence in the corresponding country shown on map. *crt:*356T was associated with emergence to artemisinin resistance mutations in a previous study in SE Asia (28).

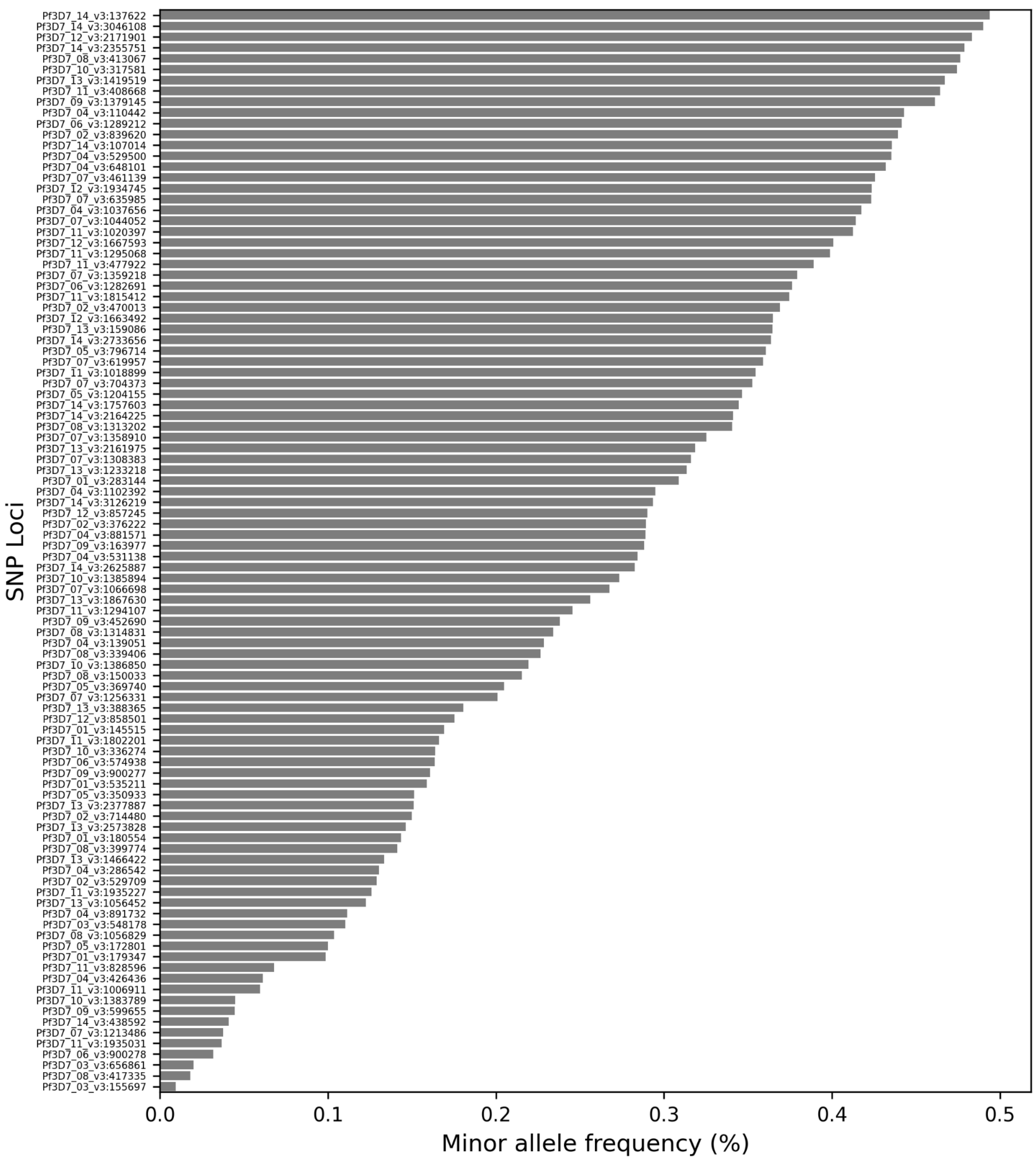


**Supplementary Figure 3. Minor allele frequency (MAF) of single nucleotide polymorphisms (SNPs) in the genetic barcodes.** Values represent the average across *P. falciparum* samples. Samples with missing calls at the respective loci were excluded from the calculations. The SNP panel was designed to capture genetic diversity across global *P. falciparum* populations. Loci are labeled as Chromosome:Position.

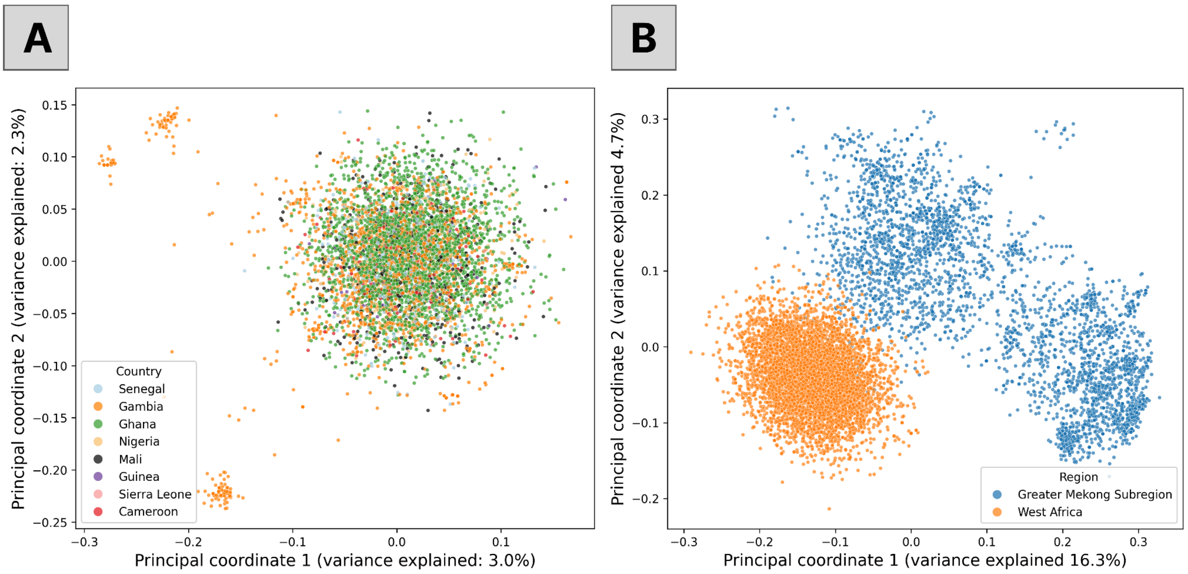


**Supplementary Figure 4. A. Principal coordinate analysis (PCoA) of 5,879 West African *P. falciparum* samples, colored by their countries.** Only samples with less than 10% missingness in their genetic barcodes and estimated complexity of infection = 1 were included. **B.** PCoA after combining the West African samples with the 5,368 *P. falciparum* samples from the Greater Mekong Subregion (GMS) public dataset (11). The first and second principal coordinates distinguish West Africa from the GMS.

**Supplementary Table 1. Prevalence of polyclonal infections across countries.**

Polyclonality is defined as a complexity of infection (COI) estimate greater than 1. The standard error (SE) is reported alongside the polyclonality estimates, and sample sizes are included in the next column.

| Country | Polyclonality (%) ±SE | No. of samples |
|---|---|---|
| The Gambia | 15.84±0.67 | 2961 |
| Sierra Leone | 20.75±3.22 | 159 |
| Senegal | 27.19±1.64 | 732 |
| Nigeria | 29.3±1.38 | 1092 |
| Mali | 31.17±1.2 | 1479 |
| West Africa avg. | 32.9±1.77 | 12393 |
| Ghana | 34.22±0.67 | 4986 |
| Guinea | 46.84±3.62 | 190 |
| Cameroon | 57.93±1.75 | 794 |

**Supplementary Table 2. Summary of *Plasmodium* species detected in samples across countries.**

| Country | *P. falciparum* | *P. malariae* | *P. ovale* | *P. vivax* |
|---|---|---|---|---|
| Cameroon | 1007 (98.63%) | 11 (1.08%) | 3 (0.29%) | 0 (0.0%) |
| Gambia | 2864 (99.97%) | 1 (0.03%) | 0 (0.0%) | 0 (0.0%) |
| Ghana | 4959 (99.46%) | 9 (0.18%) | 14 (0.28%) | 4 (0.08%) |
| Guinea | 293 (89.6%) | 25 (7.65%) | 5 (1.53%) | 4 (1.22%) |
| Mali | 1629 (98.31%) | 15 (0.91%) | 9 (0.54%) | 3 (0.18%) |
| Nigeria | 1157 (97.88%) | 17 (1.44%) | 4 (0.34%) | 4 (0.34%) |
| Senegal | 686 (98.56%) | 3 (0.43%) | 5 (0.72%) | 1 (0.14%) |
| Sierra Leone | 256 (91.1%) | 16 (5.69%) | 5 (1.78%) | 3 (1.07%) |
